# Pharmacological restoration of deficits in mitochondrial trafficking rescues aberrant axonal activity in tauopathy

**DOI:** 10.1101/2025.05.02.651902

**Authors:** Marie H Sabec, Oliver R Adam, Michael C Ashby

## Abstract

Neurodegenerative tauopathy is associated with impairments in both active axonal trafficking and presynaptic function. Although deficits in axonal trafficking reduce supply of mitochondria to presynaptic sites, where they provide a local source of energy production and calcium homeostasis, it is unclear if this contributes to progressive synaptic impairment driven by aberrant tau. Our *in vivo* two-photon imaging of mitochondrial movement and calcium dynamics in neocortical neurons of tauopathy mice revealed a progressive tau-driven decrease in axonal mitochondrial trafficking that correlates with disruption of presynaptic activity. Furthermore, we have shown that tau-driven presynaptic dysfunction can be rescued by pharmacological upregulation of axonal mitochondrial trafficking. Therefore, as well as revealing the disruptive influence of tau-induced mitochondrial trafficking impairments on presynaptic function in early stages of tauopathy, these findings highlight the therapeutic potential of targeting axonal trafficking to confer synaptic resilience to pathology.

## Introduction

The maintenance of neural and synaptic function is an energy demanding process which requires sensitive and sustained homeostatic regulation ^1^. Mitochondria located at presynaptic sites along the axonal arbor are promising candidates to both fulfil this regulatory role and provide a local energy source to support presynaptic function ^2-4^. The importance of mitochondria to synaptic and neural function is highlighted by the wide range of disorders associated with disrupted mitochondrial function ^5^. For example, deficits in mitochondrial ATP production and motility have been reported in several *in vitro* models of neurodegenerative tauopathy ^6-8^ and mitochondrial localization at synapses is altered in brains of Alzheimer’s Disease patients^9^. The active transport of mitochondria along axons is critical to deliver mitochondria, synthesized at the soma, to presynaptic sites where they can anchor or fuse with pre-existing mitochondria to support local synaptic activity ^10-13^. Pathological tau exposure induces dysregulated presynaptic activity and synaptic toxicity, leading to eventual axonal death and functional dysconnectivity^14-17^. Mitochondrial trafficking deficits present in tauopathy-affected axons could negatively impact presynaptic function and health further, but it remains unknown if restoration of trafficking can confer a resilience against the effects of aberrant tau.

To study the pathogenic impact on mitochondrial trafficking, we used the PS19 mouse model of tauopathy, which expresses the human tau gene mutation (hMAPT_(P301S)_) associated with early onset frontotemporal dementia with Parkinsonism ^14^, to quantify disease-related changes in the active transport of mitochondria, and axonal activity. Using *in vivo* two-photon imaging of cortical axons, we found a progressive impairment in axonal mitochondrial transport that coincides with altered activity patterns at presynaptic boutons in the early stages of tauopathy. Furthermore, we show that pharmacological rescue of mitochondrial trafficking deficits also restores tau-related axonal dysfunction.

## Methods

### Mice

Male and female PS19 (Prnp-MAPT*P301S hemizygous) and non-carrier littermate control mice were purchased from Jackson Laboratory (RRID:IMSR_JAX:008169) and breeding was maintained on-site. Male mice were single-housed following cranial window implantation, female mice and non-implanted male mice were group housed throughout the experimental procedure. All mice were housed under a 12hour light/dark cycle (light phase 8:00-20:00), with experimental procedures conducted during the light phase of the cycle. Water and food was available ad libitum. All procedures were conducted in accordance with the UK Animals Scientific Procedures Act (1986), in addition to local rules and regulations of the University of Bristol.

### Stereotaxic Surgeries

Surgical procedures were conducted on mice at age 8 or 20 weeks. Mice were transduced with viral vectors targeted to the right M1/2 cortex (AP; +1.3, ML; +0.7, DV; -0.3-0.7) by stereotaxic injection (StereoDrive, NeuroStar or Micro4, World Precision Instruments). Viral transduction was titred by cre-dependency to obtain sparse but high intensity expression. AAV2-hSyn-Cre-WPRE (5.048*10^10GC/ml, Addgene, #105553-AAV2) was co-injected with AAV2-Flex-GFP-P2A-MTS.STagRFP (3.22 *10^13GC/m, VectorBuilder, #VB220525-1458wse) or AAV2-Flex-axon.GCaMP8-P2A-MTS.STagRFP (3.21 *10^13 GC/ml, VectorBuilder, #VB240322-1379tuw). For in vivo two-photon imaging, mice were implanted with chronic cranial windows (3mm ∅) centred above the injection site. In brief, the overlying skull was removed with the dura left intact and a circular glass coverslip was attached with superglue and gentamicin-containing bone cement (DePuy), as previously described^37^. A circular head-bar was also affixed. Surgeries were conducted under isoflurane anaesthetic (2%, 0.6L/min) with systemic (Rimadyl; 5mg/kg, i.p. & Dexamethsone; 40mg/kg, i.p.) analgesia administered. Mice were given post-operative DietGel and monitored for 5days for weight loss or signs of pain or discomfort. Mice were excluded from the experiment if implanted windows became damaged or dislodged or if animals showed signs of ill health.

### Two-Photon Imaging

Mice were sedated (Isoflurane; 1%, 0.5L/min) and subject to imaging under a two-photon microscope at 1 month post-injection (mpi). Mice were head-fixed on a homeothermic heat pad and virally-labelled axons were imaged under a 16x (0.8NA, 3mm WD, Nikon) or 24x (1.1NA, 2mm WD, Nikon) water-immersion objective. Fluorescent signals were acquired using a resonant-galvo scanning microscope, a high performance femtosecond-pulsing laser (MaiTai, DeepSea) and ScanSci acquisition software (LabView). For mice injected with AAV-Flex-GFP-P2A-MTS.STagRFP, axons were visually identified using the GFP signal at 920nm. Fields of view (FOV; 1024 x 1024 pixels, 128μm2 or 110μm2) containing non-overlapping axons and minimal dendrites were imaged in a 5μm stack (1μm steps, 24 frames per step) to produce a high resolution structural image. Mitochondrial trafficking was then recorded in this stack at 1020nm for 3600 frames (600 stacks, 232.5sec) using a high-speed piezo giving a stack acquisition rate of 2.58Hz. For mice injected with AAV-Flex-axon.GCaMP8m-P2A-MTS.STagRFP, axons were identified using the punctate STagRFP signal from axonal mitochondria. Mitochondrial trafficking was imaged as previously described. Axonal GCaMP8 signals were then recorded at 940nm in 1.5μm steps for 3600 frames (900 stacks, 232.5sec) to give a stack acquisition rate of 3.88Hz. Isoflurane sedation and imaging was restricted to 1hour, allowing for 3-5 FOVs per animal. Mice injected with AAV-Flex-axon.GCaMP8m-P2A-MTS.STagRFP were subject to a second imaging session following Barasertib administration. Barasertib (ThermoFisher, #17140646) was dissolved in DMSO (5%), PEG300 (40%), Tween-80 (5%), and saline (50%) and injected (50mg/kg, s.c.) every other day for 2 weeks before re-imaging, with a final dose given on the day of imaging.

### Histology

Brains were extracted from naïve or 1mpi mice for tau or structural analysis, respectively. Mice were administered with non-recovery sodium pentobarbital (i.p) and transcardially perfused with phosphate buffered saline (PBS, 0.1M) and paraformaldehyde (PFA, 4%) for tissue fixation. Brains were post-fixed overnight, then cryoprotected in PBS with 30% w/v sucrose, and sectioned to 50μm by cryostat (CM3050, Leica). Slices were incubated in blocking solution for 1hour before overnight 4°C incubation with primary antibodies: Mouse monoclonal anti-DA9 (1:500, Gifted by Prof. Peter Davies, Albert Einstein College of Medicine), mouse monoclonal anti-PHF1 (1:500, Gifted by Prof. Peter Davies, Albert Einstein College of Medicine), guinea pig monoclonal anti-Bassoon (1:1000, Synaptic systems, #141.318), chicken polyclonal anti-GFP (1:1000, Abcam, #ab13970), rabbit polyclonal anti-DsRed (1:1000, Living colors, #632496). For axonal GFP amplification, with or without StagRFP amplification and bassoon staining, slices were subject to antigen retrieval in sodium citrate (pH 6.0) at 90°C for 10mins prior to blocking. Secondary antibodies were applied for 3hours at room temperature. Secondary antibodies used were anti-mouse AF488 (Invitrogen, RRID;AB_2534088), anti-chicken AF488 (Invitrogen, RRID;AB_2534096), anti-rabbit AF546 (Invitrogen, RRID;AB_2534077), and anti-guinea pig AF647 (Jackson Immuno Research, RRID; AB_2340476). For tau staining, nuclei were stained with TO-PRO-3 (Invitrogen, #T3605) before mounting. Mounted sections were cover-slipped (Prolong gold antifade mountant, Molecular probes) and imaged by confocal microscope (FV1000, Olympus). Tau images were acquired as 10μm stacks (2μm steps) by 20x water-immersion objective (0.8NA Olympus; FOV 635.9μm2, 1024×1024 pixels, av. 3 frames per step). Axon density images were acquired as 10μm stacks (2μm steps) by 25x water-immersion objective (1.05NA Olympus; FOV 253.9μm2, 1024×1024 pixels, av. 2 frames per step). Mitochondria and bassoon images were acquired as 5μm stacks (1μm steps) by 25x water-immersion objective (1.05NA Olympus; FOV 169μm2, 1024×1024 pixels). For quantitative analysis all imaging parameters including FOV size, exposure, laser power, and gain were kept consistent between samples.

### Quantification And Statistical Analysis

#### Data Analysis

All experiments were acquired and analysed blinded to genotype and conducted in a pseudo-randomized order. Mitochondrial trafficking was quantified using the “kymolyzer” plugin^42^ on ImageJ. STagRFP channel image stacks were first motion corrected using the “moco” plugin^43^. An average stack projection from the GFP channel was then used to identify <4 axons per FOV and kymographs were generated and traced from these axons using kymolyzer. Parameters of displacement distance, velocity of movement, and time spent in motion were then used to classify individual mitochondrion as motile or immobile using a two-step cluster analysis. Axonal calcium signals were extracted using the “suite2p” environment in Python^44^. GCaMP8m image stacks were motion corrected, and boutons functionally segmented before manual verification based on anatomical structure by the experimenter. Raw fluorescent traces extracted from suite2p were pre-processed for analysis by removal of 0.7*neuropil noise signal and normalisation against trace average value to give ΔF/F values. Calcium peaks were quantified from ΔF/F traces using a custom MATLAB script; In brief traces were corrected for drift using a 25-frame sliding window, and peaks exceeding 2SD were detected. The time-point for each peak was then used to calculate the bouton-bouton pairwise correlation for each bouton and FOV. Immunofluorescent images were quantified using custom ImageJ macros. For tau staining, max intensity projections were created, and soma (>15 pixels) were counted from a thresholded mask of the TO-PRO-3 channel. Tau (DA9 or PHF-1) positive cells were manually counted. For axon density, sum intensity projections were created and Li threshold masks were generated from the GFP channel, and axonal segment (>16 pixel) number and size were measured for each FOV, as previously described^45^. For mitochondria and bassoon quantification, max intensity projections were created and the GFP channel was triangle thresholded to generate a mask. Punctate mitochondrial and bassoon signals were then restricted to the axonal mask area and counted. For co-localisation mitochondrial signal was further restricted to a thresholded mask of axonal bassoon signal and counted.

#### Statistical Analysis

Data is presented as mean ±s.e.m. or median ±interquartile range depending on normality assumptions. Data was analysed by general linear model (GLM) with subject included as a random factor, unless otherwise stated. Bonferroni corrected post-hoc comparisons are included when appropriate. Data with non-normal distributions were log transformed before analysis by GLM. Population wide displacement was analysed by Kruskal-Wallis with post-hoc comparisons. The proportion of motile mitochondria was analysed by Chi Square test of proportions. The difference in within-axon vs between-axon correlation was analysed by 1-sample t-test against 0 and non-paired t-test. All statistical analysis was conducted in SPSS statistics (IBM) and graphs were created in Prism 10 (Graphpad) software, with statistical significance taken as P<0.05.

## Results

### PS19 mice have an age-dependent reduction in mitochondrial motility

To examine if axonal mitochondrial trafficking is altered in the tauopathy brain, we injected adeno-associated viruses (AAV) to express GFP and mitochondrially-targeted mito-sTagRFP in the motor cortex of hMAPT_(P301S)_ non-carrier and hemizygous (“PS19”) mice (Figure 1A). The movement of mitochondria within axons of labelled neurons were then imaged with *in vivo* two-photon microscopy in lightly anaesthetized mice aged 3 and 6 months old (Figure 1B, Supplemental Video 1). During each ∼5 minute imaging period, the majority of mitochondria displayed very little or no movement, in line with previous studies which have reported that most axonal mitochondria are stationary in adult brain ^11, 18, 19^. However, we also observed motile mitochondria with widely varying dynamics. Comparison of the 2161 mitochondria imaged across our experimental groups showed that mitochondria moved less distance in the 6-month-old compared to 3-month-old animals. This age-dependent decrease was exacerbated in the older PS19 mice, suggesting a progressive effect of the mutant tau (Figure 1C).

**Figure 1.**
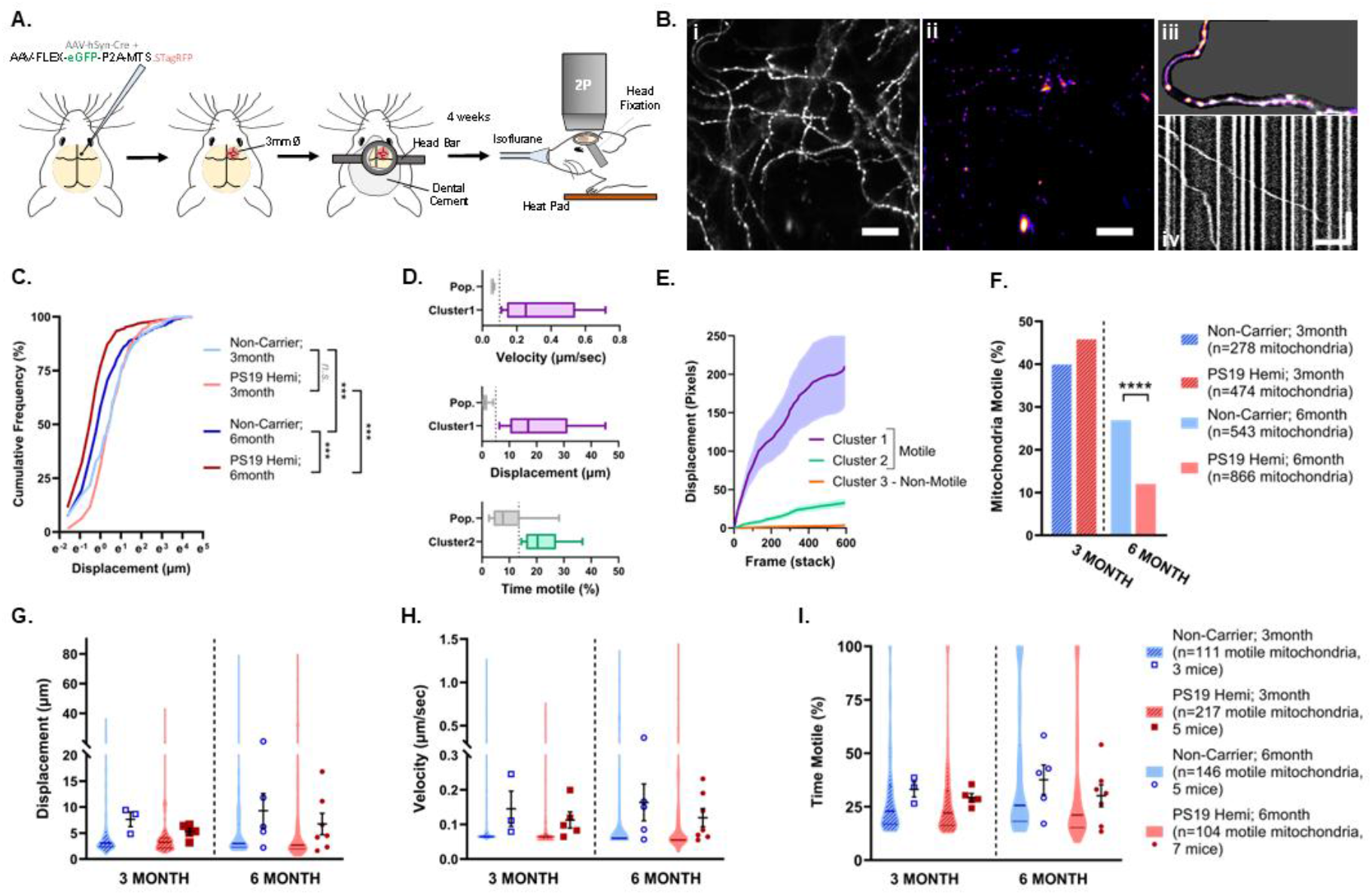
Impaired axonal mitochondrial trafficking in the brains of PS19 mice. A) Schematic representation of experimental protocol B) Representative two-photon FOV with i) axons shown in grey and ii) mitochondria in hot LUT. iii) Example axon (region indicated by white arrows in full FOV) with maximum projection of mitochondrial time-lapse overlaid and iv) corresponding kymograph showing mitochondrial paths in white with time represented in the y-axis. Scale bar = 20μm (y=60secs) C) Total distance moved by mitochondria imaged at 3 or 6 months in non-carrier and PS19 hemi mice (Kruskal-Wallis: H_(4)_=353.3, p<0.0001). Post-hoc comparisons ***p<0.001 are shown. D) Distinctive movement parameters associated with two motile subpopulations of mitochondria by cluster analysis E) Averaged displacement paths taken by a random subsample of mitochondria from the three identified clusters (n=20 mitochondria/cluster) F) Proportion of mitochondria classed as motile from non-carrier and PS19 mice (3months X^2^_(1,752)_=2.441, p=0.1182; 6months X^2^_(1, 1409)_=50.62, ****p <0.0001). G) Total distance moved by motile mitochondria (Age F_(1,13.4)_=0.212, p=0.652; Genotype F_(1,13.4)_=2.467, p=0.140; Age*Genotype F_(1,13.4)_=0.001, p=0.970) H) Average speed of movement (Age F_(1,14,2)_=0.640, p=0.444; Genotype F_(1,14.2)_=1.273, p=0.278; Age*Genotype F_(1,14.2)_=0.005, p=0.945) I) Time spent motile (Age F_(1,13.1)_=1.686, p=0.217; Genotype F_(1,13.1)_=1.993, p=0.181; Age*Genotype F_(1,13.1)_=0.147, p=0.707;). Violin plots represent the motile mitochondria population with group median ±IQR, squares/circles represent averages for individual animals with group mean ±sem.

To explore the changes driving the population-level decrease in trafficking, we classified individual mitochondria into motile and non-motile groups. A two-step cluster analysis revealed two distinct motile sub-populations. The first consisted of a small proportion of highly motile mitochondria, characterized by higher velocities during movement and greater overall distances traveled (Figure 1D; cluster 1, shown in purple). The second sub-population was distinguished by a higher proportion of total time spent in motion compared to the non-motile sub-population (Figure 1D; cluster 2, shown in green). Appropriate identification of motile and non-motile sub-populations was verified by mapping the trajectories for random subsamples (n=20 mitochondria/cluster) from each cluster, which displayed distinct profiles of movement (Figure 1E). Comparison of motile and non-motile sub-populations revealed a significant age-dependent reduction in the proportion of motile mitochondria in 6-month-old PS19 mice relative to age-matched non-carrier littermates (Figure 1F). Analysis of trafficking parameters of the motile mitochondria sub-populations did not show any significant genotype differences in terms of the total distance travelled (Figure 1G), the velocity of movement (Figure 1H), or the time spent motile (Figure 1I) at either 3 or 6 months of age. Together this indicates that tauopathy is associated with a progressive decrease in the likelihood of mitochondria becoming motile but does not impact their dynamics once they start moving.

### PS19 mice express pathological tau, but do not exhibit structural degeneration, at 6 months old

To compare tau-related pathology with the progressive decrease in mitochondrial motility, immunohistological analysis was conducted on post-mortem mouse brains. Confocal imaging of total (DA9-positive) and pathological (PHF-1-positive) tau forms in the cortex (Figure 2A) revealed an increase relative to non-carrier mice in the total tau in PS19 mice by 3 months of age, consequent to their exogenous hMAPT over-expression (Figure 2B). However, pathological tau expression was no higher than non-carrier or primary antibody lacking negative control levels at this age (Figure 2C). In contrast, by 6 months old, both total (Figure 2B) and pathological (Figure 2C) tau expression was elevated in the cortex of PS19 mice, and pathological tau levels were significantly higher than the 3-month-old PS19 mice. These findings indicate that tau overexpression alone does not reduce trafficking mitochondrial trafficking, as no deficits were observed in 3-month-old PS19 mice *in vivo*. Instead, trafficking impairments emerge alongside the development of pathological tau.

**Figure 2.**
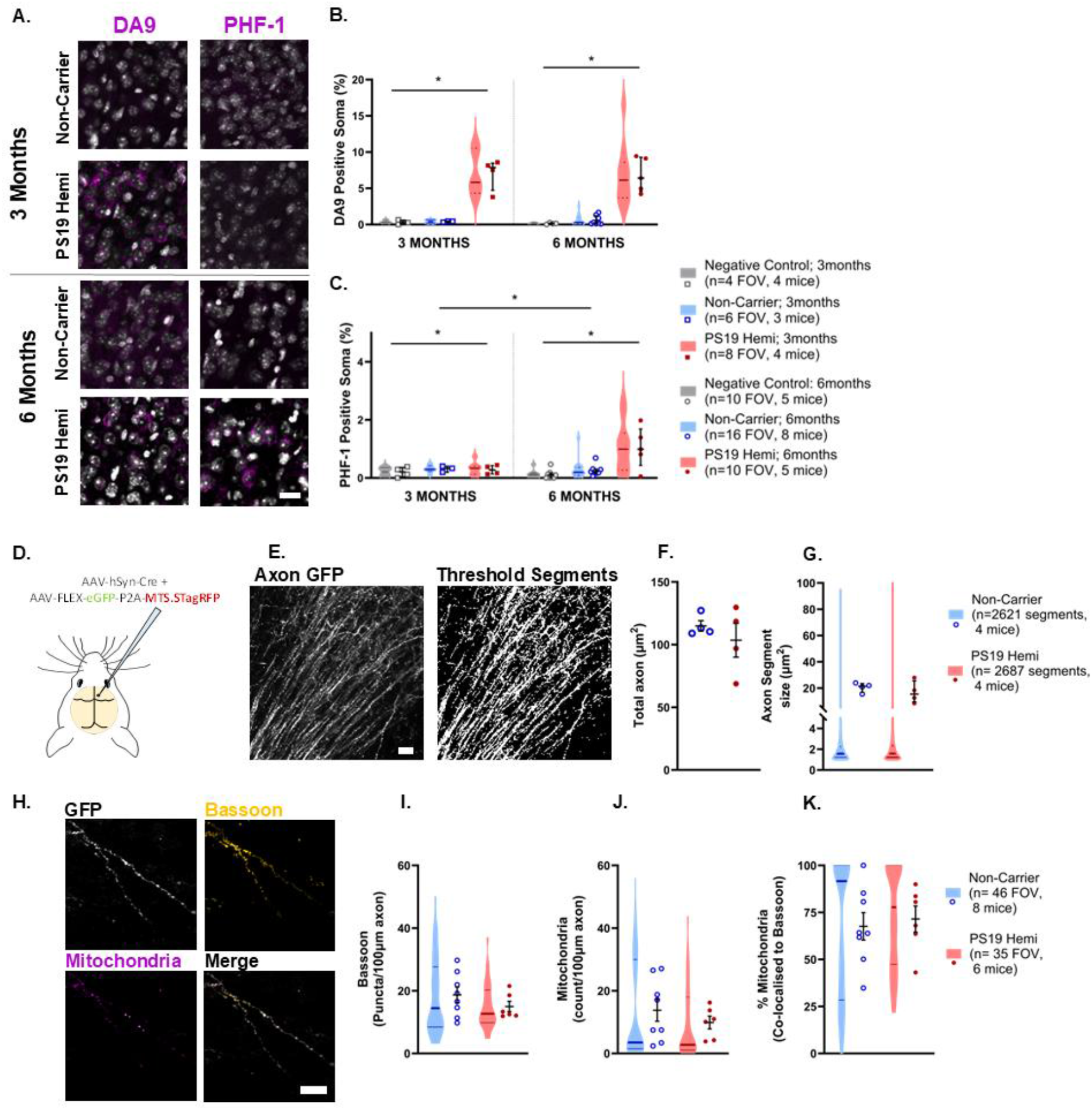
Progression of tau and axonal pathology of PS19 mouse. A) Representative ROIs showing tau (magenta) and somatic (grey) labelling from confocal images. Markers of total tau (left) and pathological tau (right) are shown in 3 (top) and 6-month-old (bottom) mice. B) Proportion of soma expressing tau in negative control (without 1° antibody), non-carrier, and PS19 hemi mice at 3 and 6 months (Age F_(1,42)_=0.007, p=0.936; Genotype F_(2,42)_=40.574, p<0.001; Age*Genotype F_(2,42)_=0.033, p=0.968). C) Proportion of soma expressing pathological tau in negative control, non-carrier, and PS19 hemi mice at 3 and 6 months (Age F_(1,32.7)_=2.090, p=0.158; Genotype F_(2,31.5)_=4.617, p=0.017; Age*Genotype F_(2,31.5)_=3.708, p=0.0.36). Violin plots represent all imaged FOVs with group median ±IQR, circles represent individual animal averages with group mean ±sem. Significant effects are shown on the graphs *p<0.05. D) Schematic representation of viral labelling technique E) Representative axon-dense FOV from confocal images (left) and thresholded mask (right) used to measure fragmentation. F) Total axonal density (t_(6)_=0.8139, p=0.4486) in masked image. G) Size of axonal fragments (F_(1,6.1)_=0.525, p=0.496). Violin plots represent all axonal segments with group median ±IQR, circles represent average fragment size for individual animals with group mean ±sem. H) Representative confocal images showing bassoon and mitochondrial labelling restricted to the GFP- positive axonal regions I) Density of bassoon puncta per 100μm length of GFP+ axon (F_(1,11.8)_=0.199, p=0.664) J) Density of mitochondria per 100μm length of axon (F_(1,11.8)_=0.456, p=0.512) K) Proportion of total mitochondrial co-localized to presynaptic bassoon puncta (F_(1,12.1)_=0.382, p=0.548). Violin plots represent all imaged FOVs with group median ±IQR, circles represent individual animal averages with group mean ±sem. Scale bars = 20μm

To investigate putative changes in axonal integrity, mitochondrial distribution, and presynaptic density, cortical AAV injections were again employed (Figure 2D). Axonal degeneration, characteristic of tauopathy, is predicted to begin with localized swelling (or “blebbing”), which precedes axonal fragmentation, leading to temporary beading and eventual cell death^20^. To identify potential blebbing or beading events, we imaged high-density regions of axonal projections near the soma using confocal microscopy (Figure 2E). No genotype differences in axonal density (Figure 2F) or fragmentation (Figure 2G) were found, nor were examples of identifiable swelling reported, suggesting that cortical axonal degeneration is not evident in our PS19 mice at 6 months of age.

We hypothesized that reductions in mitochondrial trafficking, and consequently mitochondrial supply, would also contribute to the development of synaptic deficits. To test for changes in presynaptic density and co-localization with mitochondria, we imaged immunolabelling of the pre-synaptic protein bassoon in cortical brain slices from virally transduced mice (Figure 2H). There was no difference in total density of bassoon puncta between PS19 and non-carrier mice (Supplemental Figure 1). Bassoon expression was then filtered to include only puncta expressed on virally labelled axons, but again there were no genotype-dependent differences in axonal bassoon density (Figure 2I), axonal mitochondrial density (Figure 2J) or mitochondrial co-localization to bassoon (Figure 2K). Together this indicates that whilst known pathological forms of tau are present in the PS19 mouse cortex at 6 months, this expression precedes or is not associated with significant axonal structural degeneration or synaptic loss. Furthermore, despite the observed decrease in mitochondrial motility *in vivo*, there is no reduction in mitochondrial occupancy at presynaptic sites.

### 6-month-old PS19 mice have altered axonal bouton activity

As there were no apparent structural or mitochondrial localisation deficits, we next investigated whether reduced mitochondrial trafficking may relate to changes in axonal function in 6-month-old PS19 mice. To assess this, we used axonal calcium signals as a proxy for local activity, with injection of an AAV to label mitochondria (mito-sTagRFP) and express axon-targeted GCaMP8m (axon-GCaMP8) in the same neurons (Figure 3A).

**Figure 3.**
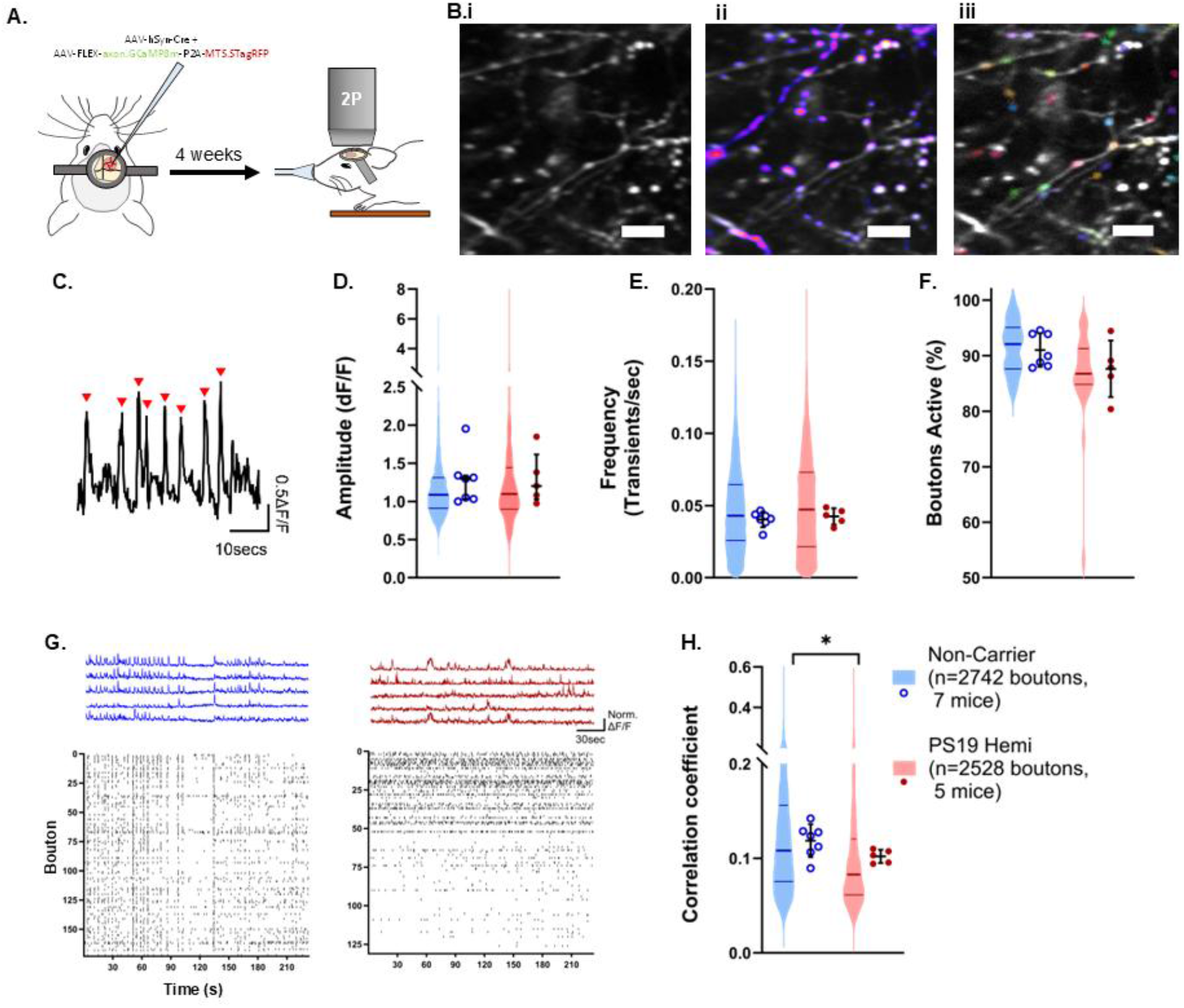
Tau-driven decorrelation in presynaptic bouton activity. A) Schematic representation of experimental protocol B) Representative portion of i) two-photon FOV with GCaMPm shown in grey, ii) the same image with the mitochondria overlaid in hot LUT showing both axonal and non-axonal mitochondria, and iii) axonal bouton segmentation shown in multi-colours following suite2p processing. Scale bars = 10μm C) Representative portion of extracted and normalised signal from an individual axonal bouton, with red arrows indicating identified calcium peaks. Scale bar =10sec (y=0.5Δ/F) D) Average calcium transient amplitudes (F_(1,9.9)_=0.111, p=0.746) E) Average calcium transient frequency (F_(1,7.6)_=0.610, p=0.459) F) Proportion of axonal boutons displaying ≥1 calcium transient during the session (F_(1,10.9)_=3.453, p=0.090) G) Example ΔF/F traces (top) and corresponding FOV raster plots (bottom) from a non-carrier (left) and PS19 hemi (right) mouse. Scale bar = 30sec (y=Norm ΔF//F) H) Pairwise bouton-bouton correlation across all FOV (F_(1,5126)_=3.973, *p=0.046). Violin plots represent all imaged boutons with group median ±IQR, circles represent individual animal averages with group mean ±sem. Significant genotype differences *p<0.05 are shown.

Once again, confirming our previous result (Figure 1), mito-sTagRFP imaging of mitochondrial dynamics showed the reduction in the proportion of motile mitochondria in the 6-month-old PS19 hemi mice relative to non-carrier littermates (Supplemental Figure 2). In the same axons, axon-GCaMP8 signals from segmented boutons were analysed to identify and characterise spontaneous calcium transients over a ∼4-minute imaging period per field of view (Figure 3B). There were no genotype-dependent differences in the amplitude (Figure 3D) or frequency (Figure 3E) of bouton calcium transients. Similarly, the proportion of active boutons (≥1 transient) was unchanged between PS19 and non-carrier mice (Figure 3F). Together, these findings indicate that, at this stage of pathology, average levels of presynaptic bouton activity across the axonal population are preserved. However, raster plots appeared to show reduced temporal synchrony of activity between individual boutons in PS19 mice compared to non-carrier controls (Figure 3G). Indeed, the pairwise bouton-to-bouton correlation coefficient across the imaging field was reduced in PS19 mice, confirming that tauopathy causes a disruption in the coordination of presynaptic activity (Figure 3H).

### Decorrelation of axonal boutons reflects impaired signal propagation within axons in PS19 mice

The overall reduction in correlation of activity between individual boutons could arise from: (1) more variability between calcium transients within the same bouton or (2) reduced coincident activity in different boutons either in the same or different axons. Abnormal mitochondrial function caused by reduced trafficking could potentially impact any of these possibilities, so we assessed each.

First, we quantified the variance of calcium transients within individual boutons to examine if disrupted mitochondrial function had led to greater within-bouton variability. There was no genotype-dependent difference in the coefficient of variation for either amplitude (Figure 4A) or area under the curve (AUC) (Figure 4B) of bouton calcium transients. This finding suggests that tauopathy does not impact the stability of calcium dynamics within individual boutons when they are activated.

**Figure 4.**
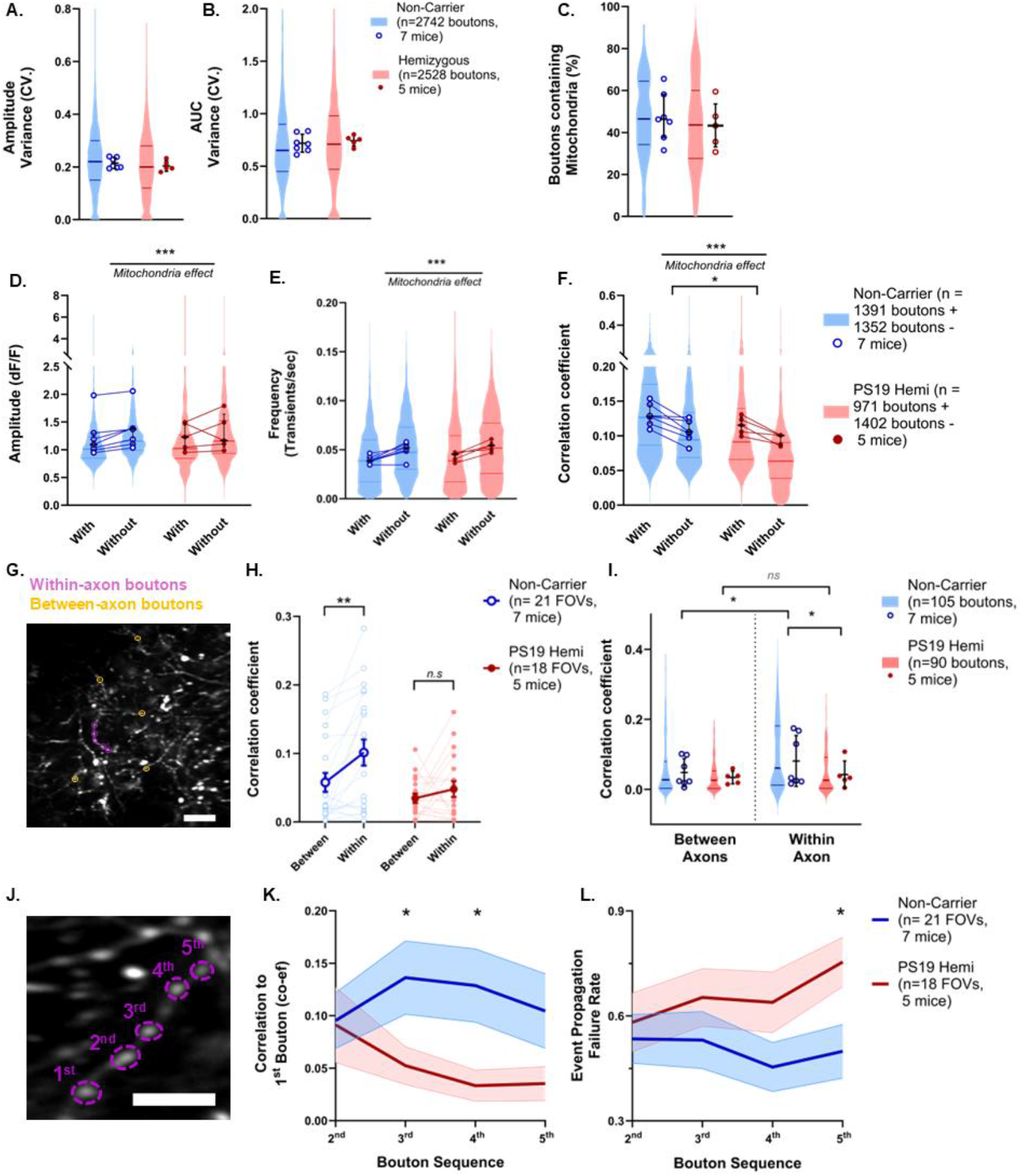
A) Intra-bouton calcium transient amplitude variance (F_(1,10.0)_=0.880, p=0.370). B) Intra-bouton calcium transient AUC variance (F_(1,10.2)_=0.417, p=0.533). Violin plots represent all imaged boutons with group median ±IQR, circles represent individual animal averages with group mean ±sem. C) Proportion of axonal boutons containing a virally-labelled mitochondrion (F_(1,9.2)_=0.004, p=0.950). Violin plots represent all FOVs with group median ±IQR, circles represent individual animal averages with group mean ±sem. D) Average calcium transient amplitudes in boutons with or without co-localised mitochondria (Mitochondria F_(1,4977)_= 178.8, p<0.001; Genotype F_(1,5111)_=0.232, p=0.630; Mitochondria*Genotype F_(1,4977)_=0.21, p=0.883;). E) Average calcium transient frequency in boutons with or without co-localised mitochondria (Mitochondria F_(1,5016)_= 184.9, p<0.001; Genotype F_(1,9.587)_=0.268, p=0.616; Mitochondria*Genotype F_(1,5016)_=0.281, p=0.596;). F) Bouton-bouton correlations across FOV in boutons with or without co-localised mitochondria (Mitochondria F_(1,4979)_= 166.8, p<0.001; Genotype F_(1,13.11)_=4.982, p=0.044; Mitochondria*Genotype F_(1,4979)_=0.254, p=0.614;). Violin plots represent all imaged boutons with group median ±IQR, circles represent individual animal averages, match across sub-population, with group mean ±sem. Significant group differences *p<0.05, ***p<0.001 are shown G) Representative GCaMP8 FOV with shown in grey and five within (magenta) and between-axon (orange) boutons highlighted. Scale bars = 20μm H) Bouton correlations for between-axon and paired within-axon boutons across the same FOV for non-carrier (t_(20)_=3.60, p=0.002) and PS19 hemi (t_(17)_=1.00, p=0.3307). Light connected circles represent bouton average difference per FOV, with group mean ±sem overlaid. Paired t-tests **p<0.01 are shown. I) Pairwise bouton-bouton correlation across boutons located on disparate axons (between) or sequentially along the same axon (within). (Grouping F_(1,376)_=22.126, p<0.001; Genotype F_(1,10)_=1.078, p=0.324; Grouping*Genotype F_(1,376)_=6.357, p=0.012;). Violin plots represent all imaged boutons with group median ±IQR, circles represent individual animal averages with group mean ±sem. Post-hoc group comparisons ***p<0.001 are shown. J) Axonal distance of sequential within-axon boutons from the first axon (Bouton Order F_(1.54,56.9)_= 207.6, p<0.0001; Genotype F_(1,37)_=0.071, p=0.791; Bouton Order*Genotype F_(1.54,56.9)_=0.304, p=0.088;). K) Bouton correlation of sequential within-axon boutons to the first axon (Bouton Order F_(2.8,92.7)_= 0.510, p=0.666; Genotype F_(1,34)_=4.753, p=0.036; Bouton Order*Genotype F_(2.73,92.7)_=1.605, p=0.197;). L) Event propagation failure rates along sequential within-axon boutons from the first axon (Bouton Order F_(2.78,103)_= 0.621, p=0.5911; Genotype F_(1,37)_=3.725, p=0.06; Bouton Order*Genotype F_(2.78,103)_=0.931, p=0.423;). Lines represent mean ±sem. Post-hoc genotype comparisons *p<0.05 are shown.

The tauopathy-driven trafficking deficits could cause changes in delivery of mitochondria to presynaptic sites and thereby impact the number of boutons that contain mitochondria and/or the replenishment of existing presynaptic mitochondria. We therefore examined whether the presynaptic bouton activity decorrelation in PS19 mice is dependent on whether boutons have resident mitochondria. On average, ∼50% of boutons in each FOV contained mitochondria, but this proportion did not differ between PS19 and non-carrier mice (Figure 4C). In agreement with our early histological findings (Figure 2H), this suggests that the decrease in mitochondrial trafficking does not impact the overall spatial distribution of presynaptic mitochondria in PS19 mice. Therefore, we investigated whether there may be a functional effect due to altered replenishment of presynaptic mitochondria by comparing calcium transients in boutons with and without resident mitochondria. Across both genotypes, boutons that contain mitochondria exhibited lower transient amplitude (Figure 4D) and frequency (Figure 4E), consistent with established roles of mitochondria in calcium buffering^13^. Reanalysis of bouton–bouton correlations within mitochondria-positive and mitochondria-negative subpopulations again revealed a significant genotype effect, with reduced coordination of activity in PS19 mice, alongside a main effect of mitochondrial content, whereby mitochondria-containing boutons exhibited higher correlation (Figure 4F). However, there was no genotype-by-mitochondria interaction, indicating that the mitochondrial support of synaptic fidelity is preserved in PS19 mice. Furthermore, it suggests that tauopathy seems to lead to presynaptic decorrelation regardless of whether boutons have resident mitochondria.

Together, these findings indicate that, despite reduced rates of mitochondrial trafficking in the PS19 mice, tauopathy does not affect the dynamics of presynaptic calcium transients, or their modulation by resident mitochondria, when they occur. Therefore, the overall reduction in bouton activity correlation likely reflects either altered co-activation across distinct neural populations within the field of view (i.e. less cell-to-cell coactivity) or impaired propagation of activity along individual axons. To distinguish between these possibilities, we sub-sampled five boutons positioned sequentially along a single axon (“within-axon”) and compared them to five boutons located on distinct, non-connected axons (“between-axon”) within the same imaging field of view (Figure 4G). As might be expected, in the non-carrier mice, the pairwise bouton-bouton correlations were higher within the same axon than between different axons. In contrast, this distinction was absent in the PS19 mice, in which within-axon correlations were similar to between-axon correlations (Figure 4H). This shows that tauopathy results in a selective reduction in the correlation of activity in boutons within the same axon (Figure 4I). There was no difference in the amplitude or frequency of activity of the within-axon compared to the between-axon boutons for either genotype confirming that the overall properties of the within- and between-axons bouton populations were comparable, distinct only in their spatiotemporal organization (Supplemental Figure 3). The fact that boutons within the same axon appear less coordinated in PS19 mice suggests that tauopathy might result in reduced fidelity of activity propagation down the axon. This leads to the hypothesis that boutons further apart would be more likely to become decorrelated by a failure of propagation. Therefore, we tested whether correlation between bouton activity is dependent on their relative axonal location, by calculating the correlation between the first bouton at one end of a chosen axon and the next four boutons moving sequentially down that axon (Figure 4J). Compared to boutons in non-carrier control mice, the sequential within-axon bouton correlation was indeed reduced in PS19 mice (Figure 4K). Furthermore, the chance of a bouton failing to show a calcium transient when the 1^st^ bouton had (failure rate) was higher in more distant boutons in PS19 mice compared to non-carrier controls (Figure 4L), consistent with a deficit in signal propagation fidelity.

Together, these findings suggest that, whilst overall presynaptic activity levels remain unaltered in trafficking-impaired axons, tauopathy drives a decorrelation among local boutons and reduced synchrony of calcium transients along sequential boutons within a given axon. This is consistent with impaired intra-axonal signal propagation rather than a global reduction in synaptic activity in the early stages of tauopathy.

### Pharmacological rescue of mitochondrial trafficking in PS19 mice also restores coordination of bouton activity

To test if reduced mitochondrial motility driven by tauopathy is directly linked to alterations in axonal activity, we used a pharmacological approach aiming to up-regulate mitochondrial trafficking *in vivo*. We used the Aurora B kinase (AuBk) inhibitor, Barasertib, which has been shown to promote neuronal mitochondrial trafficking in *in vitro* drug screens^21^. After a baseline imaging session, 6-month-old non-carrier and PS19 mice expressing mito-sTagRFP and axon-GCaMP8 were dosed with Barasertib (50mg/kg, s.c.) for two weeks and then re-imaged (Figure 5A). As in our previous experiments (Figure 1 & Supplemental Figure 2), before drug administration we observed a mitochondrial trafficking deficit in PS19 mice (Figure 5B). Following Barasertib administration, the tau-dependent reduction in the proportion of motile mitochondria was completely rescued to non-carrier levels (Figure 5B). Comparison of each mouse before and after dosing showed that Barasertib treatment consistently increased the motility ratio in PS19 mice but not in non-carrier mice (Figure 5C). In addition, Barasertib also increased the total displacement, average velocity, and time spent moving within the motile sub-population of mitochondria (Figure 5D). These results show that AuBk inhibition by Barasertib is an effective approach to up-regulate and thereby rescue tauopathy-dependent mitochondrial trafficking deficits *in vivo*.

**Figure 5.**
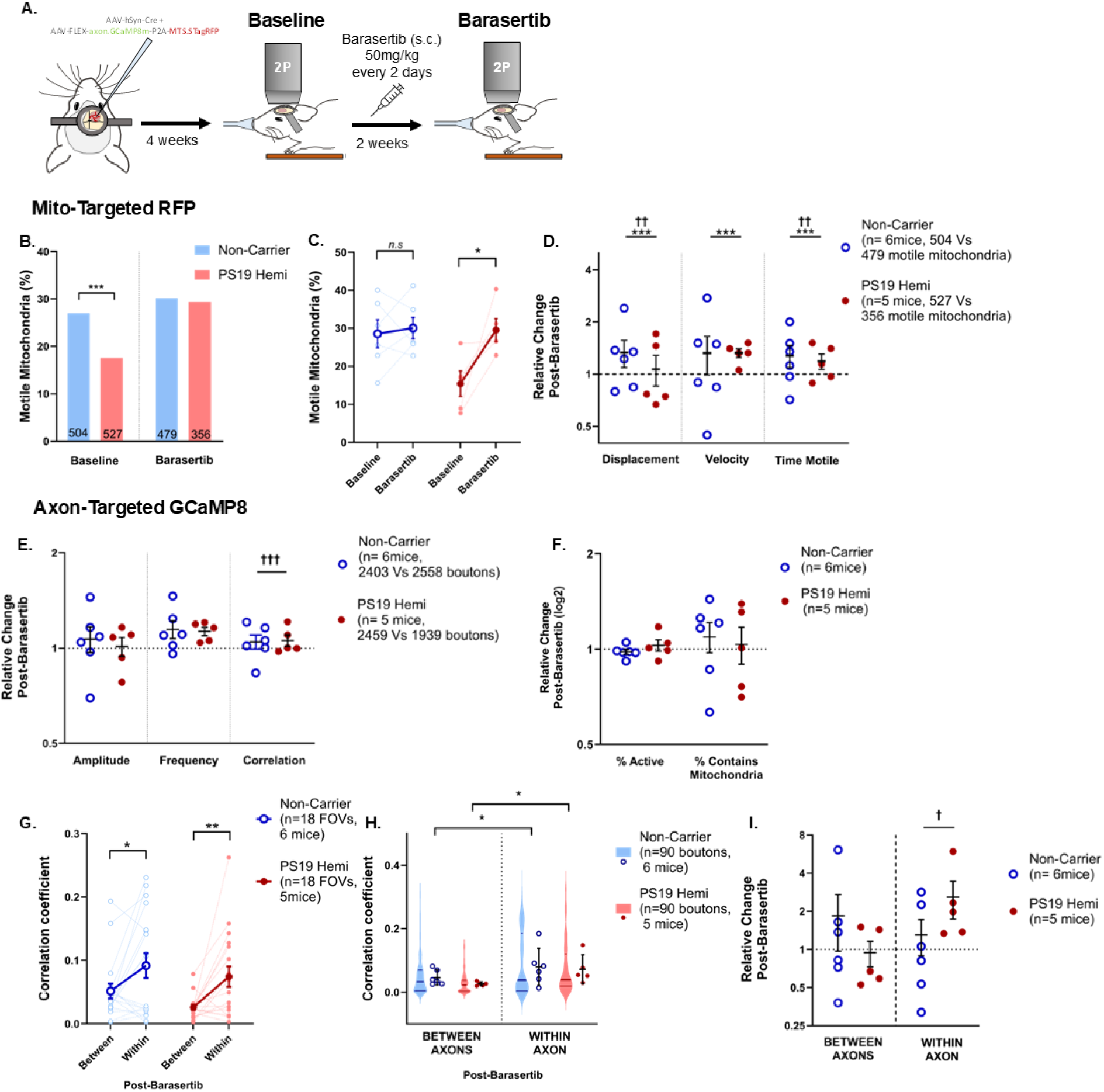
Barasertib treatment promotes mitochondrial trafficking and rescues bouton activity deficits in PS19 mice. A) Schematic representation of experimental protocol, including Barasertib dosing regimen. B) Proportion of mitochondria classed as motile from non-carrier and PS19 mice before (X^2^ _(1,1030)_=13.65, p<0.001), and following Barasertib administration (X^2^ _(1,834)_=0.060, p=0.8059). Total number of mitochondria imaged in each group/session shown at base of each bar. C) Proportion of motile mitochondria per mouse before and after Barasertib administration (Non-Carrier; t_(5)_=0.418, p=0.6933, PS19 hemi; t_(4)_=2.798, p=0.0489) D) Relative change per mouse after Barasertib administration in motile mitochondrial displacement (Drug F_(1,1856)_=14.448, p<0.001; Genotype F_(1,1911)_=0.290, p=0.590; Drug*Genotype F_(1,1856)_=7.832, p=0.005), velocity (Drug F_(1,1883)_=18.326, p<0.001; Genotype F_(1,10.4)_=2.506, p=0.143; Drug*Genotype F_(1,1883)_=1.889, p=0.169), and time motile (Drug F_(1,1877)_=11.138, p<0.001; Genotype F_(1,10.2)_=3.082, p=0.109; Drug*Genotype F_(1,1877)_=7.090, p=0.008). Significant drug effects ***p<0.001 and drug*genotype interactions ^††^p<0.01 are shown E) Relative change in bouton calcium transient amplitude (Drug F_(1,9760)_=1.713, p=0.191; Genotype F_(1,9.7)_=0.001, p=0.971; Drug*Genotype F_(1,9760)_=0.232, p=0.630), frequency (Drug F_(1,9748)_=0.947, p=0.330; Genotype F_(1,9762)_=0.059, p=0.809; Drug*Genotype F_(1,9748)_=3.483, p=0.062), and FOV-wide bouton-bouton correlation (Drug F_(1,9589)_=1.372, p=0.241; Genotype F_(1,10.3)_=3.296, p=0.099; Drug*Genotype F_(1,9589)_=17.789, p<0.001) per mouse following Barasertib administration. Significant drug*genotype interactions ^†††^p<0.001 are shown. F) Relative change the proportion of axonal boutons displaying ≥1 calcium transient (Drug F_(1,9)_=0.013, p=0.910; Genotype F_(1,9)_=1.058, p=0.331; Drug*Genotype F_(1,9)_=1.058, p=0.331) and proportion containing a virally-labelled mitochondrion (Drug F_(1,9)_=0.010, p=0.922; Genotype F_(1,9)_=1.789, p=0.214; Drug*Genotype F_(1,9)_=0.079, p=0.785) per mouse following Barasertib administration. Circles represent individual animal averages with group mean ±sem G) Bouton correlations for between-axon and paired within-axon boutons across the same FOV for non-carrier (t_(17)_=2.250, p=0.038) and PS19 hemi (t_(17)_=3.087, p=0.007) following Barasertib administration. Light connected circles represent bouton average per FOV, with group mean ±sem overlaid. Paired t-tests *p<0.05, **p<0.01 are shown. H) Pairwise bouton-bouton correlation across boutons located on disparate axons (between) or sequentially along the same axon (within). (Grouping F_(1,347)_=41.705, p<0.001; Genotype F_(1,8.9)_=0.589, p=0.463; Grouping*Genotype F_(1,347)_=0.349, p=0.555;). Violin plots represent all imaged boutons with group median ±IQR, circles represent individual animal averages with group mean ±sem. Post-hoc group comparisons *p<0.05 are shown. I) Relative change in bouton-bouton correlations across boutons located on disparate axons (Drug F_(1,367.6)_=3.880, p=0.050; Genotype F_(1,9.9)_=1.244, p=0.291; Drug*Genotype F_(1,367.6)_=0.050, p=0.823) or sequentially along the same axon (Drug F_(1,364.6)_=0.268, p=0.605; Genotype F_(1,9.9)_=0.399, p=0.542; Drug*Genotype F_(1,364.6)_=13.452, p<0.001). Significant drug* genotype interactions ^†^p<0.05 are shown.

To assess whether rescue of mitochondrial trafficking deficits impact tau-dependent changes in presynaptic activity, we also imaged spontaneous GCaMP8 fluctuations before and after Barasertib dosing in the same mice. The amplitude and frequency of axonal calcium transients (Figure 5E) and the proportion of active and mitochondria-occupied boutons (Figure 5F) were not affected by Barasertib administration in either non-carrier or PS19 mice. However, there was a significant interaction of Barasertib administration with genotype on the bouton-bouton activity correlation across each imaging FOV (Figure 5E). As Barasertib administration impacted activity correlation, we again explored if this effect was specific to changes in the correlation of boutons positioned along an individual axon or reflected a generalized increase in bouton correlation across different axons and cells. In the post-Barasertib imaging session, both non-carrier and PS19 mice exhibited significantly stronger bouton-bouton activity correlations for within-axon bouton pairs compared to between-axon bouton pairs within the same imaging field (Figure 5G&H). Furthermore, the reduced within-axon bouton correlations observed in PS19 mice during the baseline session was abolished following Barasertib treatment (Figure 5H). The rescue of within-axon bouton correlations was driven by a significant interaction of Barasertib treatment and genotype, resulting in a selective increase in within-axon correlations in the PS19 mice following drug administration (Figure 5I). In line with this, the genotype-dependent differences in pairwise bouton correlations and propagation failure rates from the first bouton across sequential within-axon bouton pairs were no longer evident following Barasertib treatment (Supplemental Figure 3). Together these effects of Barasertib indicate that pharmacological upregulation and rescue of mitochondrial trafficking can restore pathological deficits in synchrony of bouton activity within individual axons. This suggests a causal relationship between mitochondrial trafficking disruption and altered coordination of presynaptic activity in early tau pathology.

## Discussion

We have shown that axonal transport of mitochondria is reduced in the brains of PS19 mice in early stages of tauopathy. At this early stage of the disease there is not yet any clear neuronal or synaptic degeneration, but the mitochondrial trafficking deficit is linked to tau-driven decreases in coordination of presynaptic activity.

As the reduced mitochondrial motility was not associated with a decrease in mitochondrial occupancy of presynaptic boutons, we hypothesize that most axonal boutons have captured a resident mitochondrion before the onset of the pathology. The allocation of mitochondria to occupy and support new presynaptic boutons ^2^ likely contributes to the high rates of trafficking reported in developing and young neurons ^22^. Early age-dependent decreases in motility may reflect a shift in the role of motile mitochondria, transitioning from an occupancy-dominant to a replenishment-dominant function. The age-dependent decrease in mitochondrial motility between 3 and 6 months of age (Figure 1) aligns with previously reported age-dependent reductions in mitochondrial trafficking in mice up to 24 months of age ^23, 24^. Here, we have shown that the age-dependent reduction in mitochondrial trafficking is hugely exacerbated in PS19 mice as tau pathology emerges (Figure 1). This trafficking deficit is characterized by a lower proportion of motile mitochondria, without corresponding changes in the movement parameters of those that were motile. Together, these findings suggest that the reduction in motility is primarily due to impairments in the loading or unloading of mitochondrial cargo onto motor proteins, rather than deficits in motor protein dynamics. This reduction in “motile-state” mitochondria aligns well with the tau phosphatase activating domain (PAD) exposure hypothesis proposed by Kanaan and colleagues, in which conformational changes in pathological tau increase PAD exposure and consequently signal anterograde kinesin motor proteins to aberrantly detach from their membrane bound cargoes ^25-27^. An alternative mechanism by which tau has been proposed to disrupt active transport is through the destabilization of the microtubule network on which the motor proteins travel ^28, 29^. However, as axonal degeneration was not evident at the early pathological stages we studied, it is likely the decrease is due to functional rather than structural defects at this time point. As we did not observe a change in presynaptic localization of mitochondria, it is unlikely that reduced transport leads to loss of axonal mitochondria at this stage of pathology. Instead, reduced delivery of newly synthesized mitochondria may lower the rate of replacement or rejuvenation of aged mitochondria at distal axonal sites.

Selective inhibition of AuBk, including through application of Barasertib, was shown to increase mitochondrial motility in cultured cells in a small compound library screen^21^. The known canonical functions of AuBk are for cell mitosis, during which they regulate the attachment between spindle microtubules and kinetochores ^30^. Whilst recent studies have shown AuBk is also present in non-mitotic brain cells ^31, 32^, little is currently known about its function in neurons. In mitosis, AuBk can directly phosphorylate centromere-associated kinesin motor proteins to regulate their function and localization ^33^ as well as the Knl1-Mis12-Ndc80 (KMN) protein complex to modulate its microtubule binding ^34^. It is possible that neuronal AuBk has analogous effects on axonal microtubule-associated kinesin motor proteins, but this requires further investigation. We found that systemic Barasertib treatment boosted axonal mitochondrial movement dynamics in both non-carrier and PS19 mice, suggesting its enhancing mechanisms are independent of pathological tau-regulated pathways. In addition to enhancing motile mitochondrial trafficking, Barasertib also rescued the tau-dependent deficit in the proportion of motile mitochondria. Therefore, AuBk inhibition may also promote motility through tau-regulated pathways, but it is also possible that the differential increase rather reflects a ceiling effect on the maximal number of motor proteins available to transport mitochondria in 6-month-old mice. As PS19 mice had a significantly lower proportion of motile mitochondria before Barasertib treatment, they could have had more scope for a possible drug-induced increase. Increasing the pool of motile mitochondria will result in more newly synthesized mitochondria available to regenerate and/or replace aged mitochondria at presynaptic sites, potentially enhancing the resilience of synapses against a toxic tau environment ^35, 36^.

Disruption to neural activity has previously been reported in several tau models ^37^. For example, rTg4510 mice which overexpress hMAPT_(P301L)_, exhibit significant cortical hypoactivity which dominate over β-amyloid-induced hyperactivity when both pathologies are expressed ^38, 39^. We did not observe any change in activity rates in the PS19 mice, which develop pathology less aggressively than rTg4510s, but rather a bouton-bouton decorrelation of activity (Figure 3). The differences in our findings may reflect distinctions between cortical regions, the age of mice studied, or the specific tau mutation studied. A recent longitudinal study which recorded activity in mice carrying the same PS19 (hMAPT_P301S_) mutation on a human apolipoprotein E4 background likewise found no decrease in overall neuronal activity up to >9 months but did report an age-dependent decrease in the magnitude and stability of single-unit activity correlations within the hippocampus ^40^. Here, we took advantage of our optical approach to compare the coordination of activity in boutons within an axon against boutons from different neighboring axons. Changes to within-axon synchrony likely reflect a difference in axonal signal propagation, whilst changes in between-axon synchrony suggest a network level alteration of activity in local populations of cortical neurons. We found that tau primarily drives within-axon bouton decorrelation and this decrease in coordination of activity is exacerbated in boutons within the same axon that are further apart (Figure 4). As well as boosting mitochondrial transport, Barasertib treatment also rescued the tau-induced decorrelation of presynaptic bouton activity, thereby linking the two functional pathologies (Figure 5). Taken together, these findings suggest that the tauopathy-driven reductions in mitochondrial motility lead to deficits in action potential fidelity and reliable activity propagation along axons because of insufficient mitochondrial replenishment. Rather than altering bouton calcium amplitude or frequency per se, this manifests as a loss of coordinated activity across boutons within the same axon, consistent with impaired presynaptic signal transmission. Whether this deficit is specific to mitochondrial trafficking or reflects a broader disruption of axonal cargo transport, such as synaptic vesicles^41^, remains to be determined. Nevertheless, the pharmacological restoration of presynaptic activity coordination highlights mitochondria transport as a potentially tractable target for ameliorating tau-induced neuronal dysfunction.

## Data and code availability

All original code has been deposited at Zenodo at [https://doi.org/10.5281/zenodo.14779707] and is publicly available as of the date of publication.

Any additional information required to reanalyze the data reported in this paper is available from the lead contact upon request.

## Lead contact

Further information and requests for resources and reagents should be directed to and will be fulfilled by the lead contact, Dr. Michael Ashby or Dr. Marie Sabec.

## Acknowledgments

This research was funded by the Medical Research Council (grant number: MR/W005506/1). Confocal microscopy was supported by the Wolfson Bioimaging Facility at University of Bristol.

## Author Contributions

Conceptualization, M.H.S. and M.C.A.; methodology, M.H.S. and M.C.A.; Investigation, M.H.S. and O.A.; writing—original draft, M.H.S.; writing—review & editing, M.C.A.; funding acquisition, M.C.A.; resources, M.C.A.; supervision, M.H.S. and M.C.A.

## Declaration Of Interests

None.

## Supplemental Information

**Supplemental Video 1. Imaging axonal mitochondrial trafficking in the living mouse brain** Representative in vivo two-photon imaging of mitochondrial (magenta) dynamics within axonal structures (grey). The region shown is a cropped subset of a larger field of view, highlighting two motile mitochondria. Scale bar: 10 μm. Playback speed: 10× real time.

**Supplemental Figure 1.**
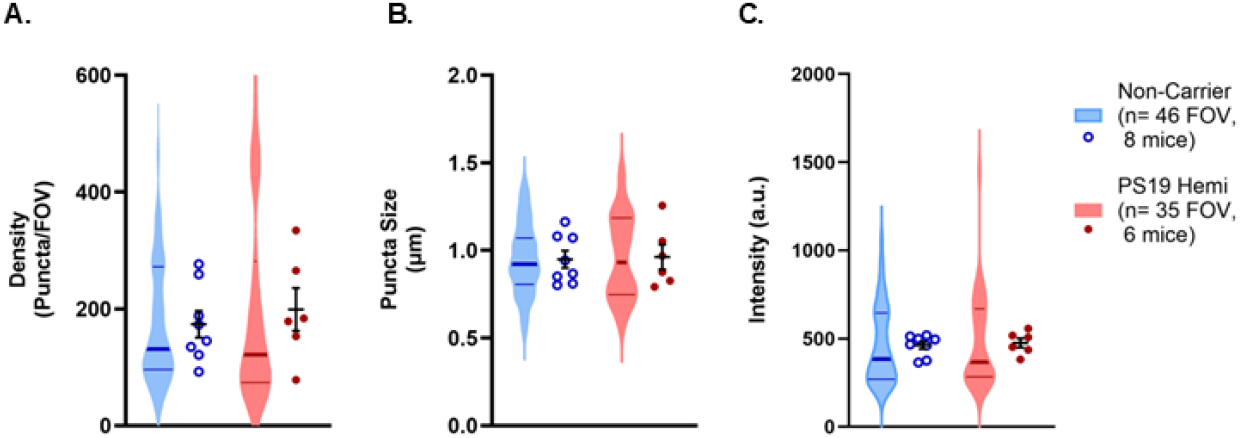
Immunofluorescent analysis of presynaptic Bassoon. A) Density of bassoon puncta detected in cortical FOV was not significantly different in non-carrier and PS19 hemi mice at 6 months (F_(11.81)_=0.409, p=0.535) B) The average size of bassoon puncta in non-carrier and PS19 hemi mice (F_(11.92)_=0.021, p=0.887) C) Average intensity of total bassoon signal detected in non-carrier and PS19 hemi mice (F_(81)_=0.031, p=0.860). Violin plots represent all imaged FOVs with group median ±IQR, circles represent individual animal averages with group mean ±sem. Significant effects are shown on the graphs *p<0.05.

**Supplemental Figure 2.**
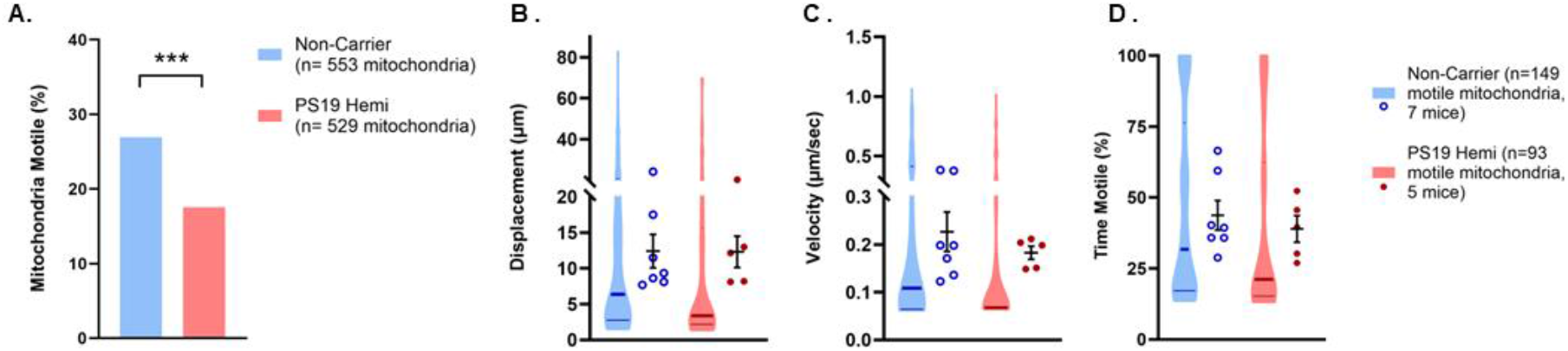
Mitochondrial tracking in axon.GCaMP8m injected mice. A) Proportion of mitochondria classed as motile from non-carrier and PS19 mice (X^2^ _(1,1082)_=13.65, p<0.001). B) Total distance moved by motile mitochondria (F_(1,11.06)_=1.234, p=0.290) C) Average speed of movement (F_(1,11.56)_=1.296, p=0.278) D) Time spent motile (F_(1,11.02)_=1.109, p=0.315).Violin plots represent the motile mitochondria population with group median ±IQR, squares/circles represent averages for individual animals with group mean ±sem

**Supplemental Figure 3.**
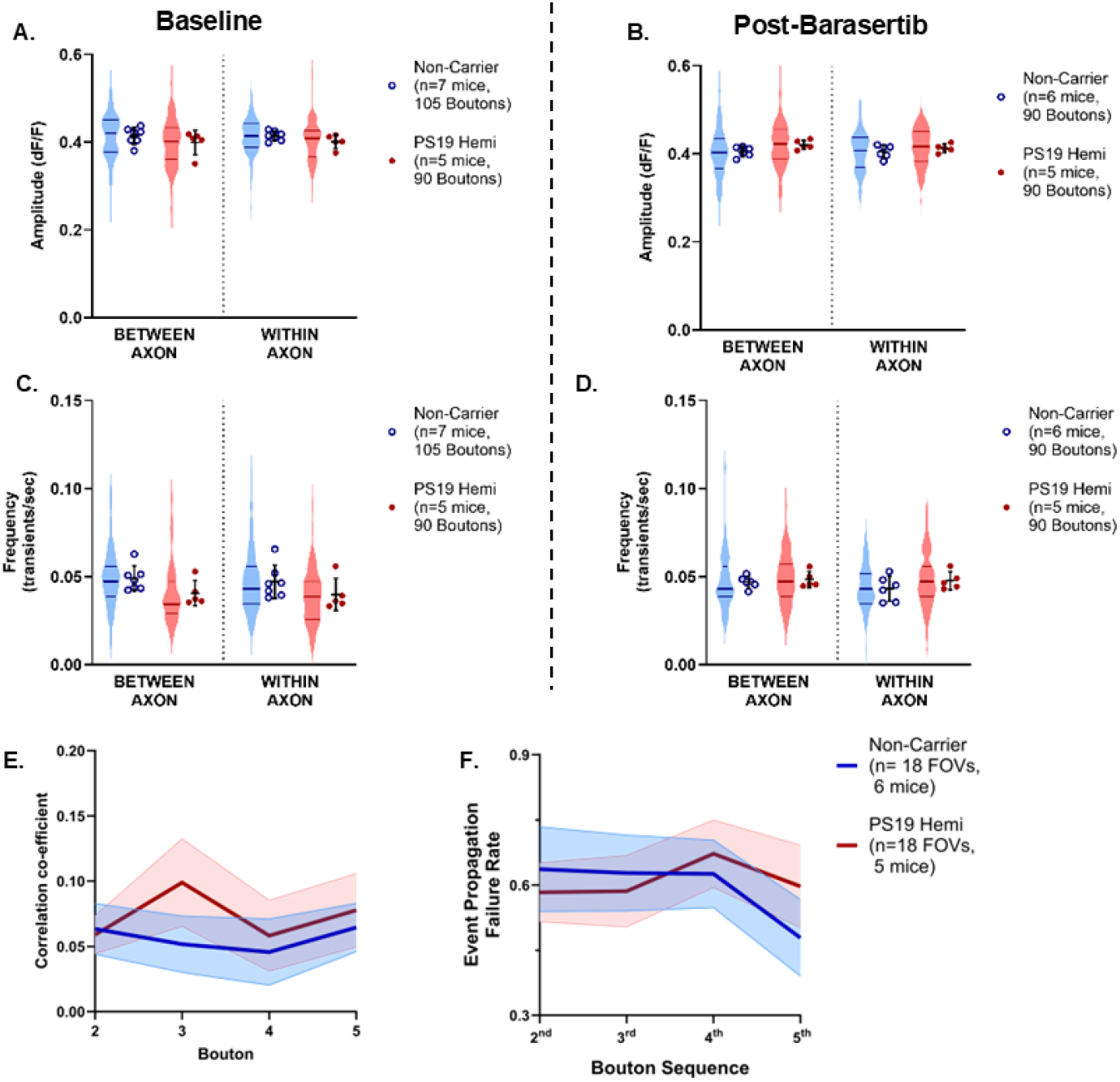
Within- and between-axon calcium transient properties. A) Calcium transient amplitude across boutons located on disparate axons (between) or sequentially along the same axon (within). (Grouping F_(1,376)_=0.371, p=0.543; Genotype F_(1,9.8)_=2.202, p=0.169; Grouping*Genotype F_(1,376)_=0.010, p=0.921;). B) Calcium transient amplitude across boutons located between or within-axon following Barasertib administration. (Grouping F_(1,348)_=0.141, p=0.708; Genotype F_(1,9.3)_=5.492, p=0.043; Grouping*Genotype F_(1,348)_=0.927, p=0.336;). C) Transient frequency in boutons between or within-axon. (Grouping F_(1,376)_=0.065, p=0.798; Genotype F_(1,10.5)_=2.715, p=0.129; Grouping*Genotype F_(1,376)_=0.001, p=0.970;). D) Transient frequency across boutons located between or within-axon following Barasertib administration. (Grouping F_(1,347)_=0.779, p=0.378; Genotype F_(1,9.8)_=1.233, p=0.295; Grouping*Genotype F_(1,347)_=0.274, p=0.601;). Violin plots represent all imaged boutons with group median ±IQR, circles represent individual animal averages with group mean ±sem. E) Bouton correlation of sequential within-axon boutons to the first axon following Barasertib administration (Bouton Order F_(2.9,97.53)_=0.514, p=0.665, p=0.666; Genotype; F_(1,34)_=0.554, p=0.462; Bouton Order*Genotype F_(3,102)_=0.552, p=0.647;). F) Event propagation failure rates along sequential within-axon boutons from the first axon (Bouton Order F_(2.59,88.0)_= 0.755, p=0.504; Genotype F_(1,34)_=0.052, p=0.821; Bouton Order*Genotype F_(2.59,88.0)_=0.572, p=0.610;). Lines represent mean ±sem

## Notes

### Competing Interest Statement

The authors have declared no competing interest.

### Summary of Updates

Manuscript revised to reflect additional data analysis and inclusion of a new results figure.

